# Modulations in motor unit discharge are related to changes in fascicle length during isometric contractions

**DOI:** 10.1101/2021.04.27.441619

**Authors:** Eduardo Martinez-Valdes, Francesco Negro, Alberto Botter, Patricio A Pincheira, Giacinto Luigi Cerone, Deborah Falla, Glen A Lichtwark, Andrew G Cresswell

## Abstract

The integration of electromyography (EMG) and ultrasound imaging has provided important information about the mechanisms of muscle activation and contraction. Unfortunately, EMG does not allow an accurate assessment of the interplay between the neural drive received by muscles, changes in fascicle length (FL) and the force/torque produced. We aimed to assess the relationship between modulations in tibialis anterior (TA) motor unit (MU) firing rate, FL and dorsiflexion torque (DT) using ultrasound-transparent high-density EMG electrodes. EMG and ultrasound images were recorded simultaneously from TA, using a 32-electrode silicon matrix, while performing isometric dorsiflexion, at diverse ankle joint positions (0° and 30° plantar flexion) and torques (20% and 40% of maximum). EMG signals were decomposed into individual MUs and changes in FL were assessed with a fascicle-tracking algorithm. MU firings were converted into a cumulative spike train (CST) that was cross-correlated with DT (CST-DT) and FL (CST-FL). High cross-correlations were found for CST-FL, 0.60 (range: 0.31-0.85) and CST-DT 0.71 (range: 0.31-0.88). Cross-correlation lags revealed that the delay between CST-FL (~75ms) was significantly smaller than CST-DT (~150ms, p<0.001). These delays affected the interpretation of MU recruitment/de-recruitment thresholds, with FL showing similar lengths for both recruitment and de-recruitment. This study is the first to demonstrate that changes in TA FL are closely related to both modulations in MU firing frequency and DT. These relationships allow assessment of the interplay between neural drive, muscle contraction and resultant torque, thereby providing a better understanding of the mechanisms responsible for the generation of muscle force.

**NEW AND NOTEWORTHY:** By employing ultrasound-transparent high-density surface EMG electrodes, we show that modulations in tibialis anterior motor unit discharge rate were closely related to both changes in its fascicle length and resultant torque. These relationships allowed quantifying delays between neural drive and muscle shortening as well as muscle shortening and torque during submaximal isometric contractions, providing an accurate estimation of the time required to generate muscle force and subsequent production of torque via the tendon.

## INTRODUCTION

One of the most fundamental issues in motor control is to understand how the nervous system interacts with muscles for the generation of appropriate joint torques and control of movement. While some answers have been obtained from separate studies in the fields of neurophysiology and biomechanics, multiple limitations from previous recording techniques have not allowed to effectively integrate these two disciplines in order to provide clearer information on how neural activity is influenced by muscle mechanics and vice-versa (19, 56). The recent integration of electromyography (EMG) recordings and ultrasound imaging has provided important information about both the mechanisms of muscle activation and contraction (2, 6, 11, 29, 48). These techniques have helped to determine for a given muscle the level of activity related to a given change in fascicle length and allowed the establishment of relationships between active/passive tissue mechanics (i.e., muscle and tendon compliance) and muscle activity. However, there are many limitations to current approaches at linking muscle mechanics with neural drive (number of motor neuron potentials received by muscles (15, 37)). For example, numerous studies have employed amplitude estimates from bipolar surface EMG recordings in order to assess changes in neural activity and the resultant force produced by muscles (28, 52, 54, 61). Due to many factors such as crosstalk, amplitude cancellation and underlying changes in muscle length and velocity, surface EMG amplitude is unfortunately poorly correlated with the resultant force produced by muscles and therefore cannot be used to directly understand the neural determinants of muscle contractions (16, 17, 42). Furthermore, studies that have examined changes in muscle architecture directly using ultrasound (3, 11, 29, 34) do not often image from the same region as where the EMG is collected (57). Therefore, changes in muscle architecture and/or morphology from nearby or other regions could potentially provide results that are not related from those studied from the region of interest. A few studies have used fine wire electromyography with ultrasound imaging to assess motor unit behaviour in relation to contraction dynamics (32, 46, 47). However, due to its high selectivity, this technique can only sample a small muscle region resulting in relatively low yields of motor units (23), limiting the ability to correlate neural drive received by muscles to the mechanical output.

Ultrasound translucent high-density EMG (HDEMG-US) electrodes have been developed to enable the simultaneous recording of large numbers of motor units and ultrasound images from the same region of muscle during contractions (5). This technique has the potential to improve our understanding of the neuromechanical determinants responsible for the generation of muscle force, however, to-date, this method has not been used to relate single motor unit discharge characteristics (e.g. blind-source separation decomposition) with fascicle movement tracked from the ultrasound images.

Our primary aim was to assess how tibialis anterior motor unit firing characteristics modulate relative to simultaneous fascicle length changes in different joint positions (short versus long muscle lengths) and at different target torques (20% and 40% of the maximum voluntary contraction, MVC). For this purpose, we examined the relationship between modulations in the motor unit cumulative spike train (CST) along with changes in fascicle length and dorsiflexion torque. We hypothesised that modulations in the CST would be strongly correlated with shortening of the fascicles, which would allow quantification of any delays between motor unit firing, muscle contraction and the torque produced via the tendon. In addition, we also examined differences in recruitment threshold in relation to fascicle length and torque, as possible delays between these signals, might affect the interpretation of motor unit recruitment and de-recruitment thresholds.

## METHODS

The study and all procedures were approved by the University of Queensland ethical committee (approval number: 2019001675) and were conducted in accordance with the Declaration of Helsinki. Ten healthy young male volunteers participated [age: mean (SD) 29 (5) years]. Exclusion criteria included any neuromuscular disorder, musculoskeletal injuries such as muscle strain as well as any current or previous history of lower limb pain/injury and be >18 or <35 years of age. Participants were asked to avoid any strenuous activity 24 hours before testing.

### Task

Participants were seated in a reclined position on the chair of an isokinetic dynamometer (Humac Norm, CSMi Computer Sports Medicine, Stoughton, USA). The right leg (dominant for all participants) was extended and positioned over a support with the knee flexed to 10° (with 0° representing full knee extension). To quantify ankle dorsiflexion torque the foot was secured to the dynamometer via non-stretch strapping tape with the approximate centre of rotation of the ankle joint (lateral malleoli) aligned to the axle of the dynamometer. Participants performed sustained and torque-varying (sinusoidal) isometric dorsiflexion contractions at different torque levels at short (ankle at 0° plantar flexion, defined as the sole of the foot being perpendicular to the shank) and long tibialis anterior muscle lengths (ankle at 30° plantar flexion). A schematic describing the setup and measurements is shown in **Figure 1**. The session began with the participants’ performing three isometric ankle dorsiflexion MVCs at each ankle angle, where each MVC was separated by 2-min of rest. The order in which the ankle was positioned (short and long tibialis anterior lengths) was randomized. Following the MVC assessment, participants were allowed to practice with visual feedback of their exerted torque (displayed on a computer monitor) performing brief ramp-hold contractions at low torque levels (20% MVC). Then, after 5 min of rest, participants performed ramp-hold (sustained) and torque-varying sinusoidal contractions at 20% or 40% MVC. For sustained isometric contractions, participants were asked to increase their torque at a rate of 10% MVC/s and then to hold the contraction at the target level for 30 s, therefore reaching the 20% MVC target in 2 s and the 40% MVC in 4 s. Two sustained 20% MVC and two sustained 40% MVC contractions were performed. For the torque-varying sinusoidal isometric contractions, the participants also reached an average torque target of 20 and 40% MVC at a rate of 10%MVC/s. However, when target torque was reached, participants had to track a sinusoidal torque target at a frequency of 0.5 Hz and amplitude modulation of 5% MVC, meaning that torque varied from 17.5% MVC to 22.5% MVC and 37.5% MVC to 42.5% MVC at 20% MVC and 40% MVC, respectively. One sinusoidal contraction per torque level was performed. These sinusoidal contractions aimed to test the ability of both the motor unit decomposition and fascicle tracking algorithms to identify motor units and track changes in fascicle length in conditions where the variability of motor unit firing and fascicle length is increased. All isometric contractions were executed in a randomized order across participants at each muscle length, but the order of the randomization was kept constant between 0° and 30° of plantar flexion.

**Figure 1.**
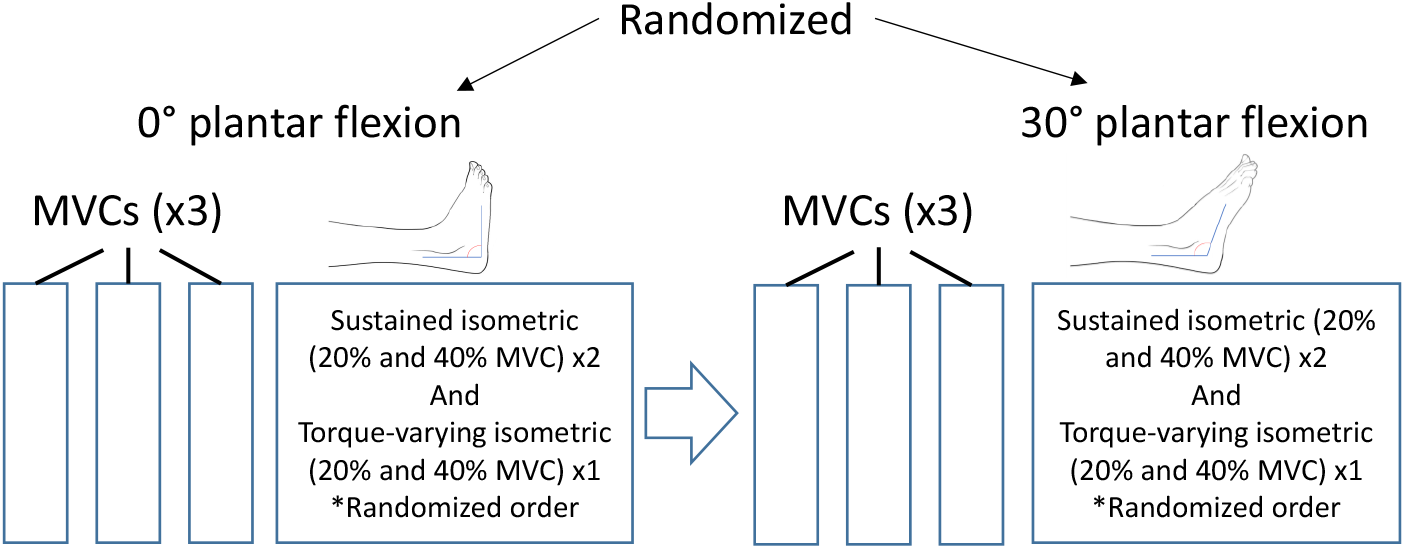
Study schematic. MVC, maximum voluntary contraction torque.

### Electromyography

Surface EMG signals were recorded from the tibialis anterior muscle using a high-density, 32-channel, HDEMG-US electrode grid (LiSIN, Torino, Italy) (5). Each grid consists of 8 x 4 electrodes (1-mm diameter, 10-mm interelectrode distance in both directions) embedded into a layer of silicon rubber (**Figure 2A)**. The array was located centrally between the proximal and distal tendons of the muscle, with the columns aligned in the direction of the fascicles as confirmed with ultrasound imaging. Skin-electrode contact was made by inserting conductive gel (Sonogel, Bad Camberg, Germany) into the electrode cavities with a mechanical pipette (Eppendorf, Hamburg, Germany) as reported previously (5). Signals were amplified and recorded using a recently developed wireless wearable HDEMG amplifier that contained a 16-bit analogue-digital converter (8) (**Figure 2B),** directly connected the electrode grids, which is ideal when HDEMG measurements are combined with other methods such as ultrasound imaging. For this experiment, data was recorded in monopolar mode at a sampling frequency of 2048 Hz, with a gain of 192 ± 1 V/V and a band-pass filter with cut-off frequencies between 10-500 Hz. Torque was also sampled at 2048Hz and recorded through the auxiliary input of the HDEMG amplifier. All EMG and torque data were processed and analysed offline using MATLAB 2019b (MathWorks Inc., Natick, MA).

**Figure 2.**
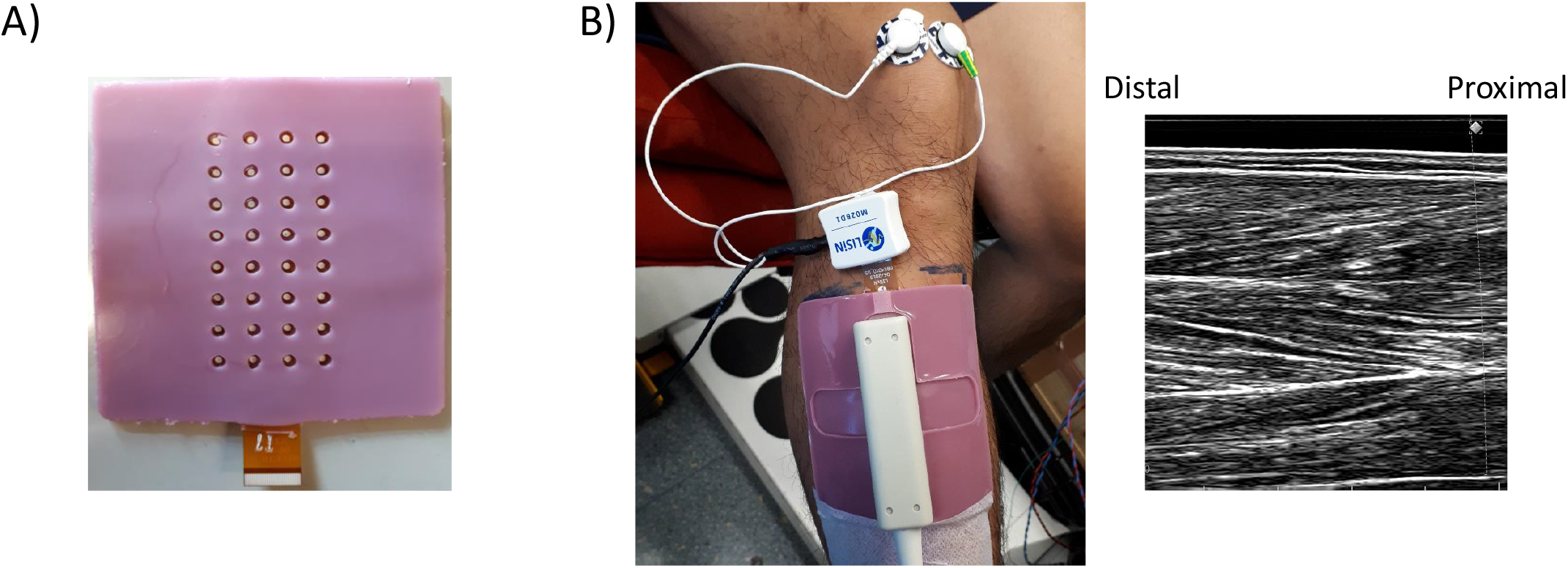
High-density surface electromyography (HDEMG) ultrasound-transparent electrodes. A) Back (up) and front (down) of the 32-channel (10 mm inter-electrode distance) electrode grid. B) HDEMG electrode grid with 32-channel HDEMG amplifier (connected on top of the electrode) and flat ultrasound probe can be seen on the left. Ultrasound image of proximal tibialis anterior muscle can be seen on the right. Note the quality of the image with accurate visualization of fascicles and, superficial, intermediate and deep aponeuroses.

### Ultrasonography

Ultrasound images of tibialis anterior fascicles were captured using B-mode ultrasonography (6 MHz, 80 frames per second, 60 mm field of view) using a 128-element multi-frequency transducer (LF9-5N60-A3; Telemed, Vilnius, Lithuania) attached to a PC-based ultrasound system (ArtUs EXT-1H scanner; Telemed, Vilnius, Lithuania). A dry HDEMG-US grid (without conductive gel) was first used to find the best alignment of the probe allowing clear visualization of the muscle’s fascicles when the probe is placed over the electrodes. Once the optimal position was identified, the skin was marked with an indelible pen. The HDEMG-US electrode grid, with conductive gel, was then positioned over the muscle at the same location and same orientation. The flat-shaped ultrasound transducer was then placed over the electrode grid and firmly strapped over the leg using an elastic bandage to prevent any movement. An example of the setup and an ultrasound image can be seen in **Figure 2B.**

### HDEMG and ultrasound synchronization

Torque, HDEMG, and ultrasound data were synchronized utilising an analogue pulse sent from an AD board (Micro 1401-3, Cambridge Electronic Design, Cambridge, UK). The trigger signal was set at 80 Hz and controlled frame by frame the ultrasound recording (i.e. every time the beamformer received a pulse, one frame of ultrasound data was recorded). The same trigger signal was sent to the auxiliary input of the HDEMG device and was aligned offline with the tracked fascicle data (see next section) obtained from the ultrasound files.

### Motor unit decomposition and fascicle length tracking

The HDEMG signals were decomposed into motor unit spike trains using an extensively validated blind source separation algorithm, which provides automatic identification of the activity of multiple single motor units (43). Each identified motor unit was assessed for decomposition accuracy with a validated metric (Silhouette, SIL), which represents the sensitivity of the decomposed spike train. Since the identification of motor unit activity with HDEMG-US grids is more challenging due to the lower selectivity of these grids (i.e., 32 channels and inter-electrode distance of 10 mm), an accuracy level of 0.86 SIL (86% of accuracy) was used to approve or discard any motor unit (usual threshold is set at 0.90 SIL (38, 43). Missing pulses producing unphysiological firing rates i.e., inter-spike intervals >250ms, were manually and iteratively included and the pulse train was re-estimated to correct its firing frequency. In cases where the algorithm incorrectly assigned two or three pulses to what was likely only a single discharge time, the operator removed this firing and the final pulse trains were reestimated as presented previously (1, 4). After this procedure, the average SIL value increased to 0.90 (0.007).

Tibialis anterior fascicle length changes were tracked offline by employing custom-made software (Farris & Lichtwark, 2016) utilizing a previously validated Lucas-Kanade optical flow algorithm with affine transformation (24, 25). From each trial, we selected the fascicle in which we were able to visualize ~80% of its length within the field of view of the image. We selected a fascicle from the anterior (superficial) compartment of the tibialis anterior muscle because 1) this was the region of motor unit activity that was likely covered by the HDEMG electrode and 2) our preliminary results showed that changes in fascicle length from this region show a crosscorrelation coefficient of at least 0.70 with the resultant torque output (see results). Changes in fascicle length were analysed according to shortening length [Δ fascicle length (i.e. difference in fascicle length from rest to target torque (20% MVC)], in order to 1) understand the amount of fascicle shortening required to recruit a motor unit and reach the required torque level (20% MVC and 40% MVC), and 2) to be able to correlate fluctuations in fascicle length with fluctuations in motor unit discharge rate, since absolute changes in fascicle length go in an opposite direction to discharge rate (fascicle length decreases when discharge rate increases, see next section). We also estimated pennation angle as the angle between the fascicle and its insertion into the central aponeurosis (50). The pennation angle was calculated when the muscle was at rest and when the torque was stable during the isometric contractions.

### Concurrent motor unit and fascicle length analysis

For sustained isometric contractions, mean discharge rate was calculated from the stable plateau torque region. Motor unit recruitment and de-recruitment thresholds were defined as the ankle dorsiflexion torques (%MVC) or fascicle shortening lengths (Δfascicle length, mm) at the times when the motor units began and stopped discharging action potentials. For these contractions we also tested the possibility to track the same motor units across the two different target torques and muscle lengths, with a previously proposed method based on cross-correlation of 2D motor unit action potentials (MUAPs) (40). In this procedure, matched MUAPs between the two trials (i.e. 20% MVC-0° vs 20% MVC-30°) were visually inspected and the two identified motor units were regarded as the same when they had a cross-correlation coefficient >0.80 (40). Cross-correlation (time domain) analysis was also used to assess the interplay between torque, fascicle length and motor unit firing data. For this purpose, motor unit discharge times were summed to generate a cumulative spike train (CST) as done previously (55). Fascicle length signals were interpolated to 2048Hz to match both motor unit and torque data. After these procedures, CST, fascicle length and torque signals were low-pass filtered (4th order zero-phase Butterworth, 2Hz) and then high-pass filtered (4th order zerophase Butterworth, 0.75Hz) as presented previously (13). These filtered signals were then crosscorrelated to assess similarities in their fluctuations (cross-correlation coefficient) and to calculate any lags between CST vs torque, CST vs fascicle length and fascicle length vs torque.

### Statistical Analysis

Unless otherwise indicated, results are expressed as mean and (SD). Normality of the data was assessed with the Shapiro-Wilk test and Sphericity was tested with the Mauchly test. Differences between fascicle parameters (absolute changes in length and pennation angle) at short and long muscle lengths during rest were assessed with paired t-tests. Absolute differences in fascicle length and pennation angle during isometric contractions were assessed with a two-way repeated measures ANOVA, with factors of muscle length (0° or 30° of plantar flexion) and torque (20 or 40% MVC). For motor unit/Δfascicle length data during isometric contractions, the following statistical tests were employed: 1) two-way repeated measures ANOVA with factors of muscle length and torque to assess differences in discharge rate 2) three-way repeated measures ANOVA with factors of muscle length, torque and signal comparison (CST vs torque, fascicle length vs CST, and torque vs fascicle length) to assess differences in cross-correlation results (correlation coefficient and lag) 3) three-way repeated measures analysis of variance (ANOVA) with factors of muscle length, torque and recruitment (recruitment vs de-recruitment) to assess differences between recruitment and de-recruitment thresholds (in terms of %MVC torque and Δfascicle length). Finally, all cross-correlation results obtained during sustained and sinusoidal contractions were averaged for each contraction type and compared by paired t-tests. All ANOVA analyses were performed using STATISTICA 12 (Statsoft, Tulsa, USA) and followed by pairwise comparisons with a Student-Newman-Keuls (SNK) post hoc test when significant. Statistical significance was set at p<0.05.

## RESULTS

### Maximal torque, motor unit decomposition and tracking during isometric contractions

The MVC dorsiflexion torque differed at the two ankle positions and was 25.2 (13.2) Nm and 51.8 (12.5) Nm at short (0° plantarflexion) and long (30° plantarflexion) muscle lengths, respectively (p<0.001). During both sustained and sinusoidal isometric contractions, an average of 7 (3), 7 (2), 7 (3) and 6 (2) motor units could be identified per participant at 20% MVC-0°, 20% MVC-30°, 40% MVC-0° and 40% MVC-30°, respectively. A representative example with the decomposition results from one participant can be seen in **Figure 3.** In this particular example, each motor unit has a clearly distinct MUAP shape (right side of the figure) which allowed accurate identification of discharge times (left side of the figure). An average of 3 (2) motor units could be tracked successfully per participant across the two muscle lengths [2D MUAP cross-correlation coefficient 0.83 (0.03)] and 3 (1) motor units per participant across the two torque levels [2D MUAP cross-correlation coefficient 0.88 (0.05)]. A representative example of the tracking procedure across the two joint angles with the HDEMG-US grids can be seen in **Figure 4**.

**Figure 3.**
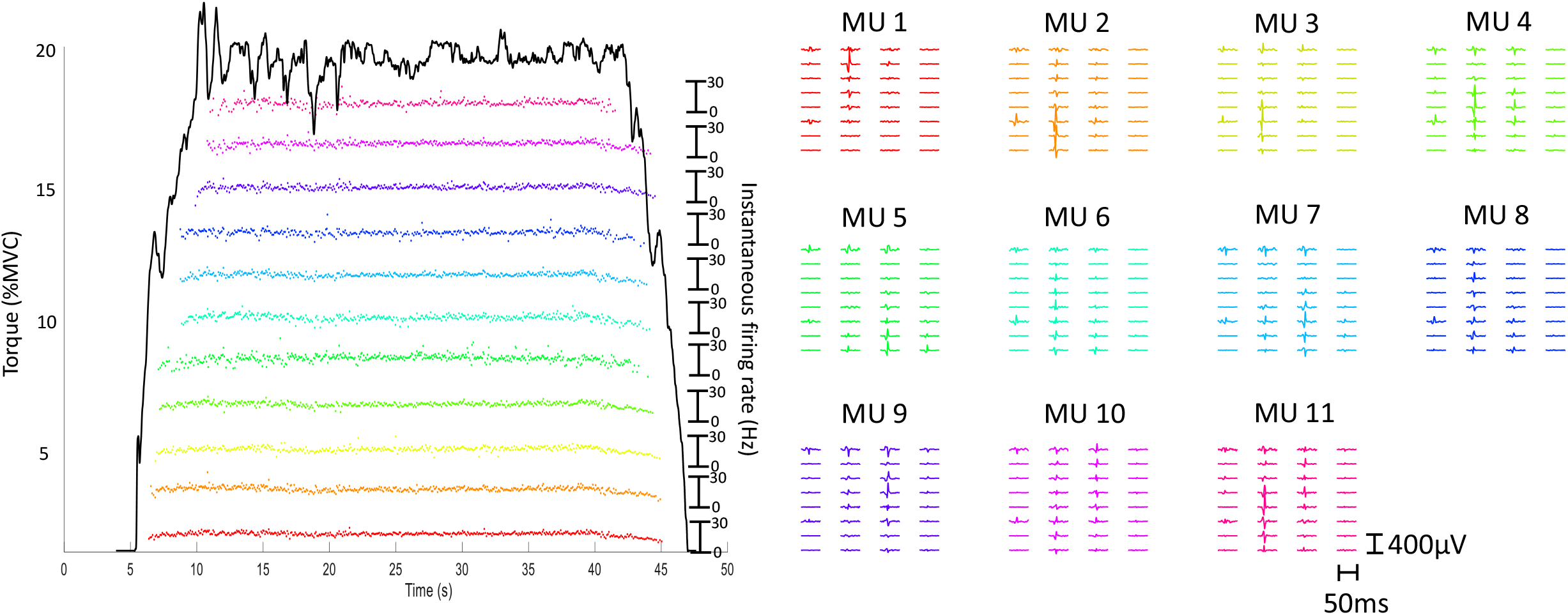
Motor unit identification during isometric contractions. A total of 11 motor units (MUs) were decomposed from the HDEMG signals in a representative participant during an isometric contraction at 20% of the maximum voluntary torque (0° of plantarflexion). Instantaneous firing rate with torque profile can be seen on the left of the figure while 2D motor unit action potentials (MUAPs) from each of these motor units can be seen on the right of the figure. Note the clear differences in MUAP shape for each of the identified units.

**Figure 4.**
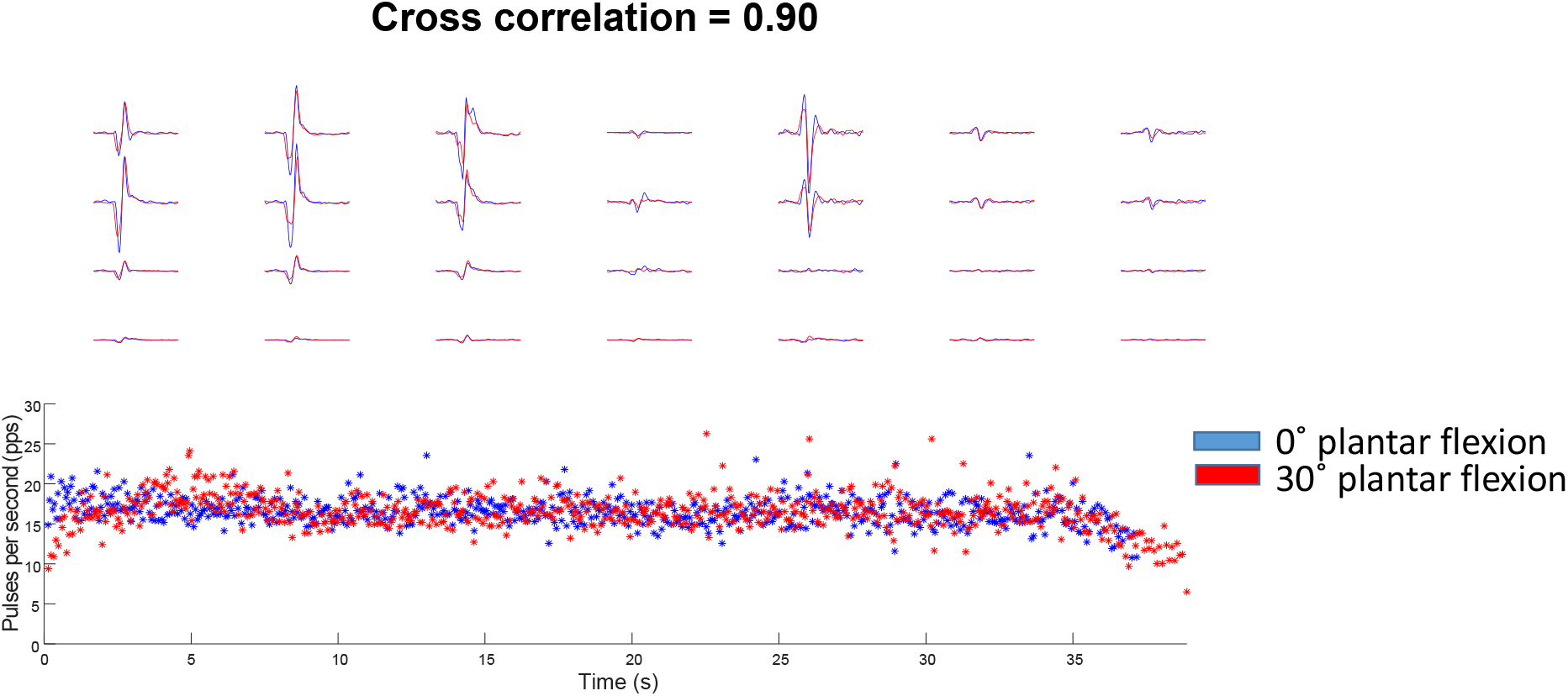
Motor unit tracking. A representative example of a motor unit that was tracked across two-plantarflexion angles at 20% MVC can be seen on the figure. For this motor unit, the action potentials (single differential) had a cross correlation coefficient of 0.90 across angles. The instantaneous firing rate of this unit can be seen on the bottom of the figure.

### Relationships between fascicle length, torque and motor unit discharge rate during isometric contractions

Absolute fascicle lengths and pennation angles during rest and isometric contractions at the two different torque targets and two joint angles are presented in **Table 1.** Overall, tibialis anterior fascicles were longer and had smaller pennation angles at 30° of plantar flexion at rest and during the contractions (p<0.001). Nevertheless, fascicle lengths decreased (torque effect: p<0.001, η^2^=0.78) and pennation angles increased (torque effect: p=0.018, η^2^=0.48) similarly with increasing dorsiflexion torque at both 0° and 30° of plantarflexion.

**Table 1.** Tibialis anterior fascicle length and pennation angles at rest and during sustained isometric dorsiflexion contractions at 20% and 40% MVC at short and long muscle lengths (different joint angles)

|  | Short (0° plantar flexion) | Long (30° plantar flexion) | P-value |
| --- | --- | --- | --- |
| Pennation angle (°), rest | 15.2 (1.7) | 12.0 (1.7) | <0.001 |
| Fascicle length (mm), rest | 60.2 (8.8) | 74.5 (9.1) | <0.001 |
| Pennation angle (°), 20% MVC | 19.2 (3.7) | 16.9 (3.9) | 0.001# |
| Fascicle length (mm), 20% MVC | 52.7 (7.5) | 66.3 (9.4) | <0.001* |
| Pennation angle (°), 40% MVC | 20.8 (3.1) | 17.4 (4.0) | 0.001# |
| Fascicle length (mm), 40% MVC | 50.8 (6.9) | 61.9 (7.9) | <0.001* |

\*Significant effect of torque ( $p < 0.001$ )
#Significant effect of torque ( $p = 0.018$ )

Similar to the changes in fascicle length, the tracking of individual motor units across short and long muscle lengths revealed similar discharge rates at the different joint angles (15.3 (2.6) Hz and 14.7 (1.4) Hz at 20% MVC, and 16.7 (2.3) Hz and 17.0 (2.4) Hz at 40% MVC, between 0° and 30° of plantar flexion, respectively, muscle length effect: p=0.74, η2=0.047). However, discharge rate increased with torque (torque effect: p=0.03, η^2^=0.43). A representative example of the concurrent assessment of changes in fascicle length, torque and CST can be seen in **Figure 5.** Briefly, after the offline identification/tracking of the fascicle (**Figure 5A**), and the identification/editing of the motor unit spike trains and subsequent conversion to CSTs, three signals were obtained: dorsiflexion torque, fascicle length and CST. All these signals were then cross-correlated on the stable torque part of the contraction (**Figure 5B**, upper panel) in order to assess common fluctuations between torque, CST and fascicle length signals. All comparisons showed high levels of cross-correlation (CST vs torque, CST vs fascicle length and fascicle vs torque, **Figure 5B**, bottom panel) and the lags obtained, showed the delay between CST vs torque, CST vs fascicle length and fascicle length vs torque. The cross-correlation results for the group of participants are presented in **Table 2**. Overall, all cross-correlation coefficients ranged from moderate to high (>0.50) showing that there was a clear relationship between CST, fascicle length and torque. It is worth noting that all correlation coefficients were higher during the sinusoidal contractions (p=0.001, **Table 2**). Nevertheless, the cross-correlation between CST vs fascicle length was significantly smaller compared to CST vs torque and fascicle length vs torque in both contraction types (signal comparison effect: p=0.007, η2=0.427 and p<0.0001, η2=0.846, for sustained and sinusoidal contractions, respectively). Cross-correlation lags (**Figure 6**) during sustained isometric contractions at 30° of plantar flexion (long muscle lengths) were larger than those at 0° plantar flexion (short muscle lengths) at both torque levels in all signal comparisons, in both sustained (muscle length effect: p=0.002, η2=0.66) and torque-varying sinusoidal (muscle length effect: p=0.034, η2=0.39) isometric contractions, with no differences between sustained vs sinusoidal isometric lag values.

**Figure 5.**
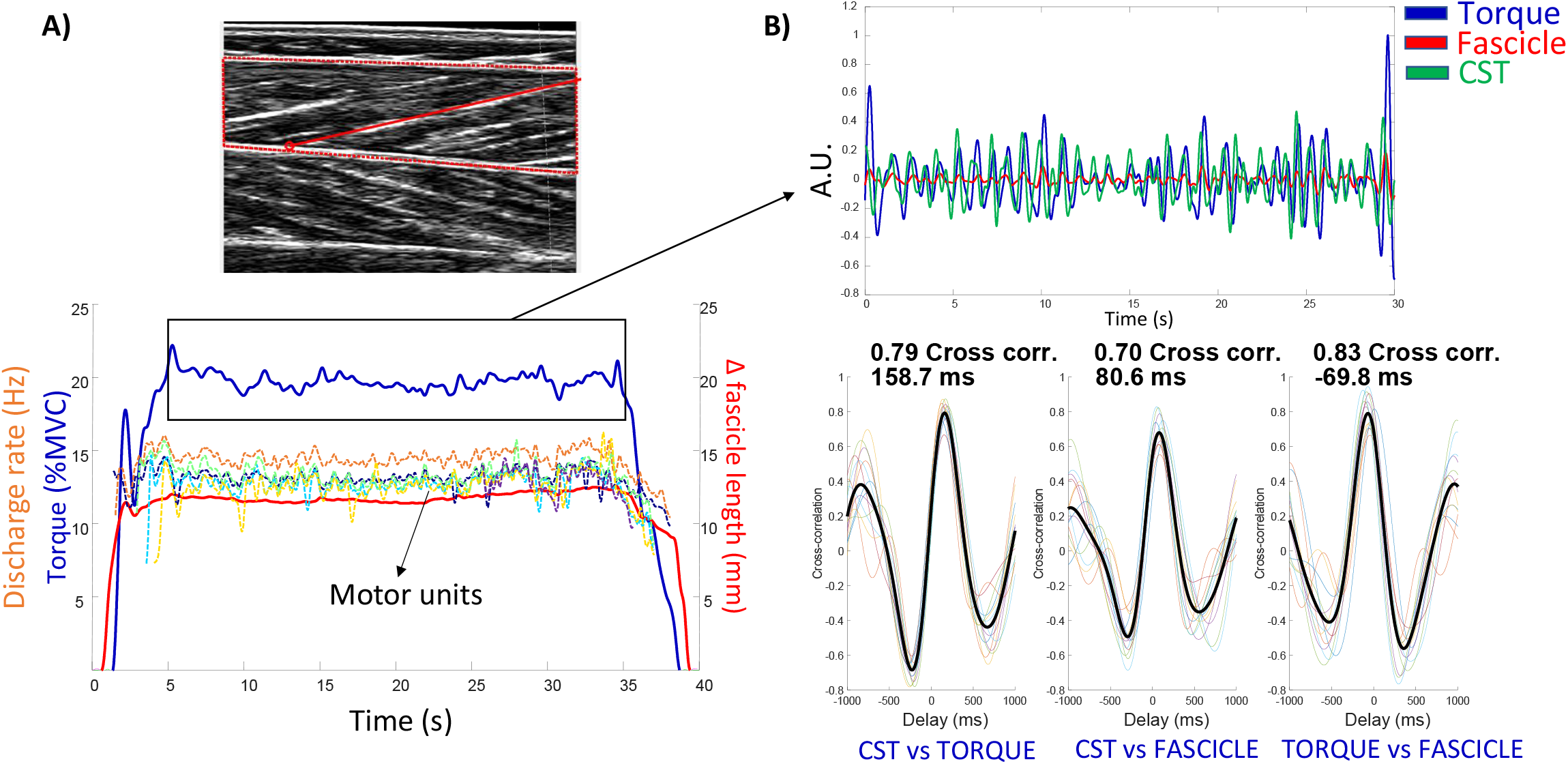
Fascicle length tracking procedure and correlation with torque and motor unit data. A) A tibialis anterior ultrasound image and a fascicle of interest (red) can be seen on top of the figure. The length data obtained from the tracking of this fascicle was then correlated with torque and cumulative spike train (CST) signals (bottom). Fascicle length data is presented as the amount of shortening from rest to target torque (fascicle length during rest minus fascicle length reached at target torque). Note that fascicle shortening precedes the generation of torque during the ramp-up phase of the contraction and then returns to baseline values after the torque signal returns to zero. The steady torque part that was selected to correlate signals is inside the rectangle. B) Common fluctuations from the three generated signals (torque, CST and fascicle length) in the steady-torque part of the contraction can be seen on top of the figure. Cross-correlation and lag (delay, ms) results between CST vs torque, CST vs fascicle length (fascicle) and fascicle vs torque can be seen on the bottom of the figure.

**Figure 6.**
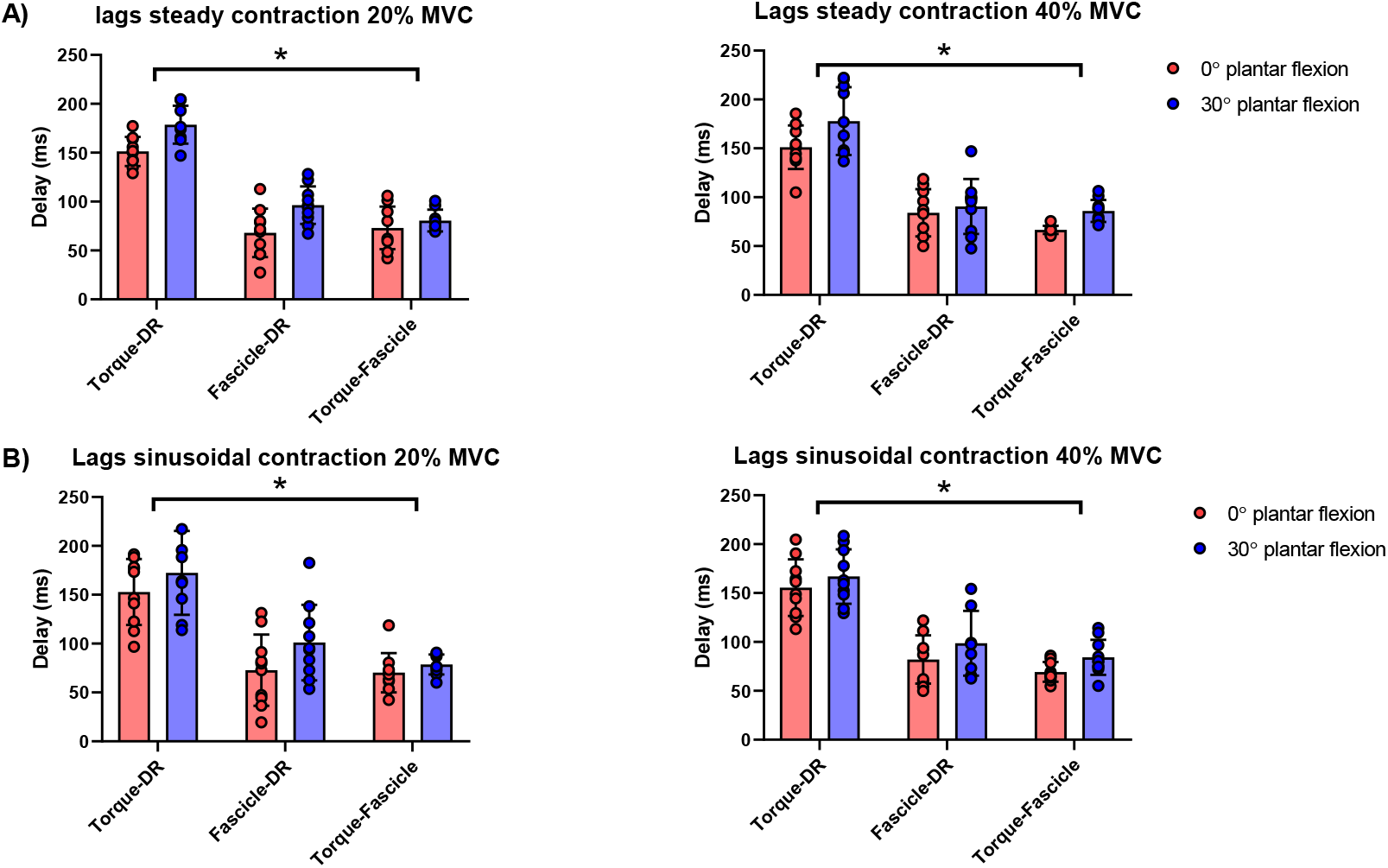
Cross-correlation lag (delay) results during sustained and sinusoidal isometric contractions. Delays between cumulative spike train (CST) vs torque, CST vs fascicle length (fascicle) and fascicle vs torque can be seen for sustained (A) and sinusoidal (B) isometric contractions at 0° and 30° of plantarflexion at 20% MVC and 40% MVC. *, significant effect of joint angle (p<0.05).

**Table 2.** Cross-correlation coefficients for comparisons between cumulative spike train (CST) vs torque, CST vs fascicle length and torque vs fascicle length during isometric contractions.

|  | Short length<br>20% MVC | Long Length<br>20% MVC | Short length<br>40%MVC | Long length<br>40% MVC |
| --- | --- | --- | --- | --- |
| Steady isometric contractions |  |  |  |  |
| Torque vs CST | 0.74 (0.65-0.88) | 0.74 (0.62-0.85) | 0.74 (0.63-0.85) | 0.61 (0.31-0.78) |
| Fascicle vs CST | 0.64 (0.51-0.81) | 0.57 (0.31-0.75) | 0.63 (0.33-0.85) | 0.50 (0.31-0.66) |
| Torque vs Fascicle | 0.71 (0.40-0.88) | 0.67 (0.33-0.89) | 0.78 (0.33-0.93) | 0.72 (0.50-0.91) |
| Oscillatory isometric contractions |  |  |  |  |
| Torque vs CST | 0.78 (0.66-0.89) | 0.77 (0.62-0.83) | 0.77 (0.66-0.89) | 0.66 (0.45-0.82) |
| Fascicle vs CST | 0.71 (0.58-0.86) | 0.73 (0.69-0.78) | 0.74 (0.60-0.86) | 0.62 (0.41-0.75) |
| Torque vs Fascicle | 0.87 (0.75-0.95) | 0.89 (0.75-0.95) | 0.91 (0.80-0.96) | 0.89 (0.77-0.96) |
| Results are expressed as mean (min-max range). |  |  |  |  |

### variations in tibialis anterior fascicle length and recruitment/de-recruitment thresholds during isometric contractions

Motor unit recruitment and de-recruitment thresholds in terms of both Δfascicle length (mm) and torque (%MVC) are shown for a representative participant in **Figure 7A**. The figure shows that recruitment threshold torque is higher than the torque at de-recruitment, however, the Δfascicle length at which motor units were recruited and de-recruited was similar. This was confirmed in motor units that were tracked across short and long muscle lengths as derecruitment thresholds were consistently lower when assessed in terms of %MVC torque (recruitment-de-recruitment effect: p=0.029, η^2^=0.43, **Figure 7B**) but similar in terms of Δfascicle length (recruitment-de-recruitment effect p=0.805, η^2^=0.007, **Figure 7B**,). In addition, although recruitment and de-recruitment thresholds increased in terms %MVC torque across 20% and 40% MVC levels (torque effect: p=0.001, η^2^=0.701), motor units were recruited and de-recruited at a similar Δfascicle length with increasing torque (torque effect: p=0.731, η^2^=0.014). Finally, recruitment-de-recruitment thresholds also increased at long muscle lengths (0° plantar flexion vs 30° plantar flexion) when considered as %MVC torque (muscle length effect: p<0.001, η^2^=0.77) but not in terms of Δfascicle length (muscle length effect: p=0.389, η^2^=0.014).

**Figure 7.**
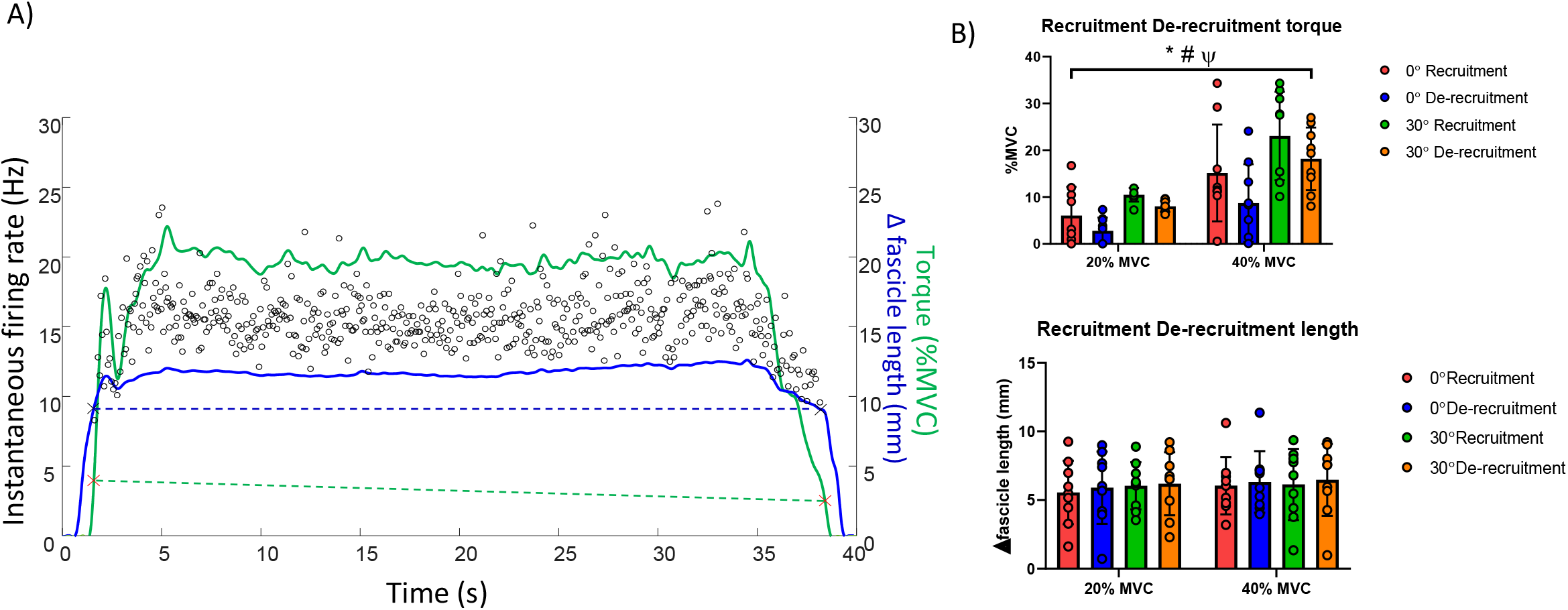
Recruitment and de-recruitment thresholds in relation to torque and fascicle length. A representative example of recruitment and de-recruitment threshold of a motor unit in relation to torque and fascicle length can be seen on the left of the figure. The recruitment threshold for this unit was higher than the de-recruitment threshold when calculated as %MVC torque (green dashed line) but similar when calculated as fascicle shortening length (blue dashed line). Fascicle length data is presented as the amount of shortening from rest to target torque (fascicle length during rest-fascicle length reached at target torque). The same results can be appreciated by the group of participants on the right of the figure as recruitment thresholds are consistently higher than de-recruitment thresholds across target torques and angles when considered as %MVC torque (upper right) but similar when calculated from fascicle length data (lower right). *, significant effect of recruitment-de-recruitment (p<0.05). #, significant effect of joint angle (p<0.05). Ψ, significant effect of torque (p<0.05).

## DISCUSSION

This is the first study showing that modulations in motor unit discharge rate are closely related to both changes in fascicle length and joint torque during isometric contractions. These relationships allowed us to quantify the delays between motor unit firing activity and fascicle shortening and, fascicle shortening and joint torque during voluntary contractions. By employing this analysis, we demonstrated that the delays between the neural drive and fascicle length or the generated force/torque are larger when the muscle contracts isometrically at longer fascicle lengths (**Figure 6**). In addition, due to the possibility to track motor units across different muscle lengths and target torques, we were also able to show that recruitment and de-recruitment thresholds are similar when considered in terms of fascicle shortening length, meaning that motor units are recruited and de-recruited at the same change in fascicle length, regardless of the joint position or torque exerted. Taken together, the examination of these relationships allows for an accurate assessment of the conversion of neural activity into muscle contractions.

### CST, fascicle length and torque relationships

A number of studies have shown that fluctuations in firing rate are closely related to the fluctuations in torque/force (14, 42, 55). Considering this observation, we attempted to quantify the level of correlation between fluctuations in firing rate (CST), changes in fascicle length and torque, in order to first, understand whether the information obtained from the identified motor units was linked to the fascicles of the region of interest (CST vs fascicle length correlation) and also to confirm that motor unit firings identified with HDEMG-US grids would be able to predict the fluctuations in torque produced via the tendon. The findings showed that all of the signals compared had moderate to high levels of correlation, confirming that the motor unit data provided a good representation of the changes in fascicle length of the region of interest. Moreover, all correlation levels increased further when greater torque variability was induced during sinusoidal contractions, which shows that motor unit activity is modulated together with changes in fascicle length, providing further confirmation of the close relationship between motor unit discharge and fascicle length. It is important to mention that this greater variability did not affect cross-correlation lags as similar delays were observed in both sustained and oscillatory contractions (**Figure 6**). Recent studies correlating both motor unit firings and torque have shown that the delay between the CST and torque decreases at higher contraction velocities (14, 36). It is likely that the frequency at which the participants matched the sinusoidal target (0.5Hz) was similar to the involuntary torque modulations induced during sustained isometric contractions. Indeed, both Del Vecchio et al. (14) and Martinez-Valdes et al. (36) found even larger delays between CST and torque (~300ms) when the contractions were modulated at slow frequencies (0.5 and 0.25 Hz respectively). Nevertheless, it is likely that the differences in delays between the present research and previous studies is mainly related to variations in the measurement apparatus (i.e., force quantified via a strain gauge versus a torque dynamometer) (9). This provides an advantage in using ultrasound techniques to assess changes in neural activity in relation to muscle contractile dynamics, as these measurements are not affected by differences between dynamometers and could also be used to isolate individual muscle dynamics for joints where multiple muscles actuate the joint.

The lower correlation between CST and fascicle length (**Table 2**) during both sustained and sinusoidal isometric contractions could be explained by a number of factors, including the subtle inaccuracies in fascicle length determination from ultrasound imaging and potential mismatches between fibres contributing to CST and the fibres in the imaging region. It is possible that the fascicle tracking or the sampling frequency employed to obtain the images (80Hz) was not sensitive enough to detect small changes in fascicle length, particularly during sustained contractions. However, the sampling frequency is well above the frequency of variation in force during isometric contraction (<5Hz (21)) and tremor (5-13Hz, (60)). Therefore, it is more likely that limitations in imaging the muscle in a single plane and the resolution of the tracking algorithm may contribute to the lower correlations. The depth at which we tracked the fascicles (superficial portion of the tibialis anterior) vs region of motor units recorded may also contribute to discrepancies in the signal correlations; the HDEMG-US electrodes are most likely to only detect motor unit activity of the most superficial motor units (20). If we also consider that blind-source separation motor unit decomposition techniques favour the identification of motor units showing the largest MUAPs and the highest spatial localization on the HDEMG grid (30, 43), it is very likely that the motor units included in the analysis mainly represent the superficial region of tibialis anterior. Nevertheless, it is important to note that we obtained moderate-high cross-correlations between CST and fascicle length during isometric contractions, which demonstrates feasibility of using such approach to understand the link between motor unit firing properties and contraction dynamics.

### CST, fascicle length and torque delays

One of the most interesting findings from this study was the possibility of quantifying delays between CST vs. fascicle length and fascicle length vs. torque, during active contractions, which can provide a better understanding of the neuromechanical interactions during movement. Historically, previous studies aiming to quantify the time required to convert neural activity into mechanical output, calculated the time-difference between the onset of force/torque and muscle activation (7) during both voluntary and electrically-induced contractions. This assessment is commonly known as the electromechanical delay, and has provided important information about the time required for a muscle to generate force/torque from a passive state. Since this analysis commonly neglects the lag between the generation of a contraction and the transmission of force to the tendon and then transmission to the measuring apparatus (which generates great variability in results (59)), more recent studies have aimed to quantify these lags by combining conventional bipolar EMG or HDEMG amplitude and ultrasound (3, 58). Similar to the current study, this previous research also revealed that fascicle shortening happens before force/torque is measured (3, 34, 58). However, the magnitude of these delays is considerably smaller than the ones presented herein, with ~6ms (EMG-FL) and ~12ms (EMG-force) of delay for electrically-induced contractions (34, 44) and ~29ms (EMG-FL) and ~50ms (EMG-force) of delay for voluntary contractions (3). Besides the multiple methodological limitations of the aforementioned approaches such as issues with the correct detection of the onset of muscle activation from EMG amplitude (18, 31) and weak correlations between EMG amplitude and force (42), it is difficult to transfer the information obtained from these delays to functional contractions. Moreover, estimates of the electromechanical delay are calculated at a single time-instant when the muscle activates from a passive state, neglecting the influence of the muscle’s afferent sensors (i.e. muscle spindle, golgi tendon organ), active motor unit twitch properties and active/passive muscle tissue interactions responsible for the regulation of muscle force during contractions (10). On the contrary, the cross-correlation of signals obtained from HDEMG-US motor unit decomposition and ultrasound could provide a better estimation of these delays, as such assessment considers mechanisms responsible for contraction dynamics (i.e. fascicle length responses to firing rate modulations) rather than just assessing delay differences between signals at contraction onset.

The longer delays reported in our study are likely due to twitch characteristics of the active motor units (14) and changes in MUAP duration during the contraction. Indeed, a very recent study from Cudicio et al. (10) which employed a new technique of motor unit twitch estimation based on deconvolution of torque and motor unit spike trains reported values of motor unit twitch time-to-peak (~110ms) similar to the delays between CST and torque reported in our study. Moreover, the delays reported in the present investigation between CST and fascicle length (~75ms) are in line with those reported in a previous study from Hamner and Delp (27), where the authors calculated the lag between EMG and simulated muscle activations (when muscle force is produced) during running at 5 m/s. We can therefore speculate that the delays quantified between CST and fascicle length have the potential to explain changes in force transmission during the execution of more functional tasks. However, this would require additional testing of the proposed approach during dynamic contractions.

### CST, fascicle length and torque delays at different joint positions

Another interesting result obtained from the cross-correlation lags was that both CST vs. fascicle length and fascicle length vs. torque delays increased when the muscle contracted isometrically in a lengthened position (**Figure 6**). Changes in the duration of muscle-fibre twitch force and muscle-tendon compliance at larger muscle lengths likely influenced these delays. To provide support to the first observation, several studies have reported that muscle twitches are significantly larger (greater contraction and half-relaxation times) when the muscle is lengthened (10, 35, 53). Therefore, it is very likely that an increase in the overall duration of the motor unit responses at longer lengths is the reason for the more delayed muscle contraction and transmission of force to the tendon observed at the two submaximal normalized torque targets (20 and 40% MVC). In addition, changes in tendon and aponeurosis elasticity during lengthening could also reduce the amount of neural drive required to activate the muscle (33, 41, 49) and potentially increase the delays between signals. Nevertheless, the assessment of both motor unit contractile properties and the effect of passive structural properties of the muscle on motor unit behaviour needs to be confirmed in future investigations. The assessment of these delays holds great potential in both healthy populations (e.g. aging) and patients (e.g. cerebral palsy), as they could help to quantify impairments in force transmission, when muscles have altered architecture (e.g. muscle contractures or sarcopenia).

### Recruitment vs. De-recruitment threshold discrepancies

The discrepancy between %MVC vs fascicle length recruitment and de-recruitment thresholds was an interesting observation. Motor unit studies typically use torque/force to assume changes in muscle behaviour, however, delays between motor unit firing and torque due to force transmission delays (along muscle/tendon and also within dynamometer) could create offsets when comparing recruitment (torque rise) vs decruitment (torque drop) thresholds. The findings from this study suggest that it is important to consider these delays when interpreting recruitment and de-recruitment results from torque/force signals since we were able to observe that recruitment and de-recruitment thresholds differed when quantified as % MVC torque (recruitment threshold was higher than the de-recruitment threshold) but were similar when quantified in terms of fascicle length. To our knowledge, this is the first study showing this discrepancy. Moreover, we were able to observe that the length at which a motor unit was recruited and de-recruited was maintained across joint angles and torque levels. Fascicle shortening preceded torque generation (as expected), but fascicles returned to resting values after the dorsiflexion torque returned to 0 Nm. This means that during the ramp-down phase of the contraction, there is a time-point where the muscle returns passively to its original length (**Figure 7**). This can explain why many studies show lower torque values for de-recruitment than recruitment when considered as %MVC torque or absolute torque (12, 39, 51) and emphasizes the need to interpret recruitment vs de-recruitment relationships with caution, when quantified from the torque produced at the dynamometer. Interestingly, similar divergences were recently reported when considering recruitment and de-recruitment in terms of joint angle or fascicle length during isometric plantarflexion contractions at varying knee-joint angles (32).

### Limitations and future developments

In this study we were able identify an average of 7 motor units across contractions, which is lower than the number of units identified with non-transparent HDEMG electrodes, where approximately 15 to 20 motor units can be identified on the tibialis anterior muscle on average per participant (39). These differences can be attributed to a number of factors. First, the grid of electrodes employed in the current study contained 32 electrodes vs. the 64 electrodes conventionally employed to decompose EMG signals. It has been shown that a larger number of channels enhances the spatial identification of MUAPs and therefore improves the separation of the multiple motor unit sources from the HDEMG with blind-source separation methods (22, 43). Second, the inter-electrode distance of the HDEMG-US grid (10mm) is less selective than the 64-channel grids (8mm) commonly employed to decompose motor unit activity, which can again influence the separation of MUAPs from the HDEMG. Therefore, improvements in electrode construction will likely increase the number of motor units identified with HDEMG-US. Nevertheless, it is important to mention that the cross-correlation between motor unit discharge rate and torque was high (~0.70) and similar to the values reported in previous studies (14, 42, 55). As mentioned previously, muscle fascicle imaging using ultrasound also has some limitations in terms of the resolution of the image limiting the tracking of small changes in fascicle length, as well as the limitation of only being able to image one plane of the muscle. Improvements in ultrasound probe construction and/or the addition of another ultrasound probe in a different portion of the electrode grid would likely help to improve estimations of changes in fascicle length during isometric contractions. Finally, it is worth mentioning that the accurate assessment of motor unit and fascicle data across torque levels and joint positions, opens up the opportunity to explore more diverse isometric contraction models (i.e., ballistic contractions or faster sinusoidal contractions) and shortening and lengthening contractions. Our present results were not affected by more dynamic variations in isometric torque and the tracking of motor units across joint angles revealed that MUAPs did not change substantially within the range of motion investigated (**Figure 4**). Very recent studies employing HDEMG recordings (without ultrasound) have successfully identified motor units during shortening and lengthening contractions (26, 45), therefore, it is very likely that future studies assessing changes in motor unit activity and fascicle length will be able to assess these interactions during dynamic conditions.

## Conclusion

In conclusion, this study shows that modulations in motor unit firing rate are closely related to changes in fascicle length during isometric contractions. The determination of these relationships allows quantifying delays between MUAP propagation, muscle contraction and subsequent transmission of force to the tendon during sustained and functional torque modulation, providing a better representation of the mechanisms responsible for the control of muscle force during active contractions.

## ACKNOWLEDGEMENTS

This study was partially funded with the University of Birmingham International Engagement Fund (BIEF) awarded to E.M.-V.

## DISCLOSURES

No conflicts of interest, financial or otherwise, are declared by the authors.

## Author contributions

Experiments were performed at the School of Human Movement and Nutrition Sciences, University of Queensland, Queensland, Australia. E.M.-V. designed research; E.M.-V and P.P. performed experiments; E.M.-V. and F.N. analysed the data; E.M.-V., F.N., A.B., G.C., D.F., P.P., G.L. and A.C. interpreted the data and contributed to the drafting of the article. All authors have read and approved final submission.

## Notes

### Competing Interest Statement

The authors have declared no competing interest.

